# Sensorimotor faculties bias perceptual decision-making

**DOI:** 10.1101/2024.05.14.594024

**Authors:** Jan Kubanek, Lawrence H Snyder, Richard A Abrams

## Abstract

Decision-making is a deliberate process that seemingly evolves under our own volition. Yet, research on embodied cognition has demonstrated that higher-order cognitive processes may be influenced, in unexpected ways, by properties of motor and sensory systems. Here we tested whether and how simple decisions are influenced by handedness and by asymmetries in the auditory system. Right- and left-handed participants performed an auditory decision task. In the task, subjects decided whether they heard more click sounds in the right ear or in the left ear, and pressed a key with either their right or left index finger, according to an instructed stimulus-key assignment (congruent or reversed). On some trials, there was no stimulus and subjects could choose either of the responses freely. When subjects chose freely, their choices were substantially governed by their handedness: Left-handed subjects were significantly biased to make the leftward choice, whereas right-handed subjects showed a substantial rightward bias. When the choice was governed by the sensory stimulus, subjects showed a rightward choice bias under the congruent key assignment, but this effect reversed to a leftward choice bias under the reversed key assignment. This result indicates a bias towards deciding that there were more clicks presented to the right ear. Together, our findings demonstrate that human choices can be considerably influenced by properties of motor and sensory systems.

## 1. Introduction

Decision-making is a hallmark of higher-order cognition. When we make a decision, we weigh the evidence that supports each alternative and choose the alternative that appears to be associated with a better outcome. This deliberate process appears to function under our own volition, independently of particular properties of the systems that provide the input to or the output from this process.

However, contrary to this impression, neuroscientists have recently suggested that variables related to simple perceptual decisions are detectable in motor circuits (Gold and Shadlen 2000, 2007; Selen et al. 2012; Kubanek et al. 2013). This opens the possibility that perceptual decisions may be *influenced* by particular properties and asymmetries of motor (Hicks and Kinsbourne 1976; Corballis 1997; Bryden et al. 1994; Bishop et al. 1996; Gabbard et al. 1998; Calvert 1998; Stins et al. 2001) and sensory (Kimura 1961a,b; Broadbent and Gregory 1964; Knox and Kimura 1970; Kimura 2011) systems.

This article investigates the influence of two of such asymmetries—handedness within the motor domain, and a right ear advantage within the sensory domain.

Handedness influences and may in part be defined by a subject’s choice of which hand to use to perform a complex movement, such as a reach for an object (Hicks and Kinsbourne 1976; Corballis 1997; Bryden et al. 1994; Bishop et al. 1996; Gabbard et al. 1998; Calvert 1998; Stins et al. 2001). We asked whether handedness could also affects perceptual decisions that are communicated using simple movements that do not require dexterity.

The auditory system exhibits asymmetries in discrimination tasks based on auditory evidence. In dichotic listening tasks, subjects discriminate verbal or non-verbal auditory stimuli that are simultaneously presented to both ears. It has been found that when the sounds are of a verbal nature, subjects typically show a “right-ear advantage” (Kimura 1961a,b; Broadbent and Gregory 1964; Kimura 2011). When the sounds are non-verbal, subjects tend to show a left-ear advantage (Knox and Kimura 1970; King and Kimura 1972; Kimura 2011). In light of these effects, we investigated whether the auditory system shows an asymmetry in a perceptual choice task that requires an accumulation of auditory evidence over a brief period of time. We specifically used a choice task in which the individual quanta of auditory evidence followed in a brief succession such that they could not be counted, and in which the auditory quanta were trivial and did not carry semantic information (clicks sounds; in contrast to words (Kimura 1961a,b; Broadbent and Gregory 1964; Kimura 2011)). This simple paradigm minimizes the involvement of higher order cognitive processes beyond those required for a decision, such as counting or word recognition.

## 2. Methods

### Subjects

Fifty-four Washington University undergraduate students (37 females, 17 males), aged 18 to 21 (mean 19.2) participated in this study. All subjects were healthy, had normal hearing capacity, and gave an informed consent. Subjects participated for class credit.

### 2.2. Apparatus and procedure

Subjects sat in a comfortable chair 70 cm in front of a flat-screen monitor. Subjects wore headphones (MDR-V600, Sony), which presented a stereo auditory stimulus (see Auditory stimulus). The volume in the left channel was set to the same level as the volume in the right channel. The subjects’ hands were comfortably positioned at a computer keyboard with the left index finger placed over the left Alt key and with their right index finger placed over the right Alt key. The control of the experimental design was accomplished using a custom program written in Matlab (The Mathworks, Inc., Natick, MA).

Each trial started with the presentation of a red fixation cross, 2 degrees in size. Subjects were instructed to fixate at the center of the cross. At the same time, subjects were presented with a stereo auditory stimulus (click sounds, see Auditory stimulus), 1.0 s in duration (Fig. 1). After the stimulus had been presented, the fixation cross shrank to 1 degree and changed its color to green. This event cued the subjects to make a movement (choice). Subjects performed 2 blocks of 300 trials each, with a brief break in between. In the first block of 300 trials, subjects were instructed to press the left Alt key with their left index finger if they heard more clicks in the left ear and to press the right Alt key with their right index finger if they heard more clicks in the right ear. In the second block of 300 trials, this instructed key assignment was reversed. The first block was completed by all 54 subjects, the second block by all but one subject.

**Fig. 1.**
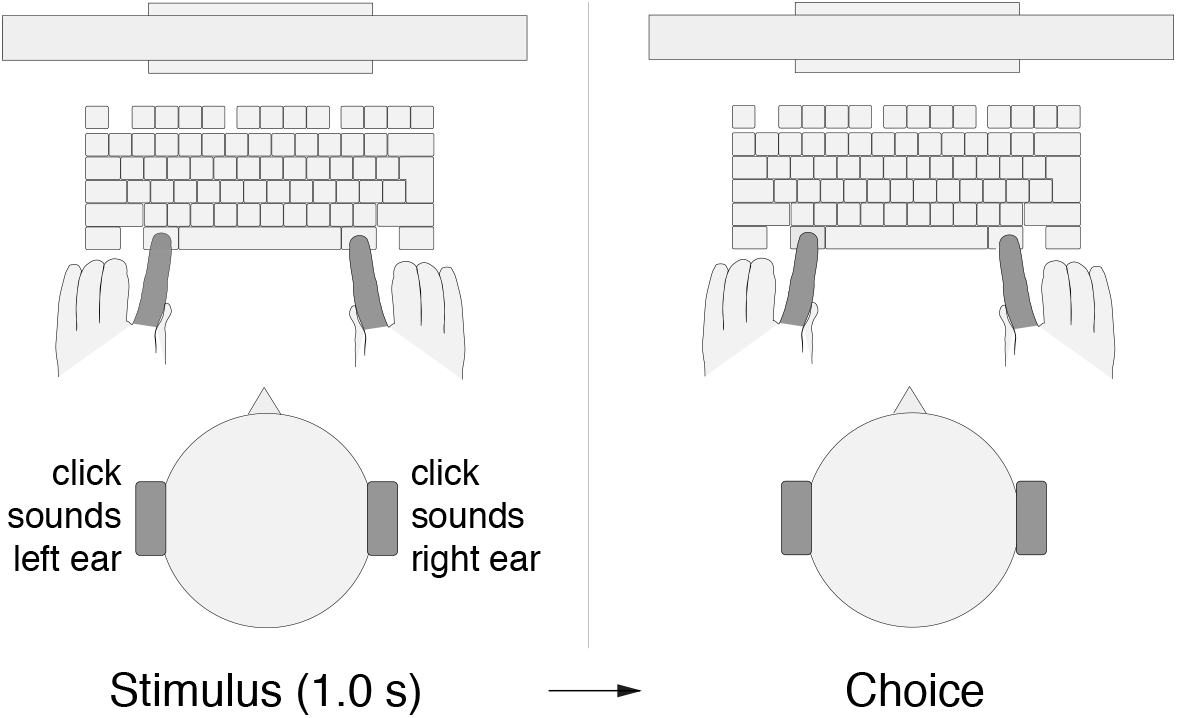
Perceptual decision task. Subjects listened to a binaurally presented auditory stimulus that comprised a 1.0 s train of Poisson-distributed click sounds (Methods). Following the stimulus presentation, subjects pressed either the left Alt key with their left index finger or the right Alt key with their right index finger, depending on a particular key assignment.

On 20% of the trials (randomly selected), no auditory stimulus was presented. When no sound was heard, subjects were instructed to choose either key (i.e., to either press the left key with the left index finger or the right key with the right index finger). The purpose of these trials was to study choice that is self-initiated by the subject.

If subjects responded prior to the green go cue or if they failed to indicate a response within 1200 ms after the go cue, the trial was considered invalid, and was aborted and excluded from the analyses. The type of error was indicated to the subjects in red, large-font text (‘TOO EARLY’, ‘TOO LATE’). Overall, the proportion of valid responses was 95.6 ± 6.1% (mean±s.d.) in the first block, and 96.4 ± 13.6% in the second block. A response was immediately followed by a display of a feedback. Specifically, a correct response was followed by the display of a green string that was randomly drawn from the set {+5*c*, +10*c*, +15*c*, +20*c*, +25*c*}. An incorrect response was followed by the display of a red string randomly drawn from the set {−5*c*, −10*c*, −15*c*, −20*c*, −25*c}*. The feedback was displayed for 0.5 s. The next trial started immediately following the offset of the feedback.

### 2.3. Auditory stimulus

The auditory stimulus presented to each ear consisted of a train of brief (0.2 ms) click sounds drawn from a homogeneous Poisson process (Kubanek et al. 2013). Each train lasted 1.0 s. The stereo stimulus was composed such that the number of clicks presented to the left ear (*C*_*l*_) plus the number of clicks presented to the right ear (*C*_*r*_) summed to a fixed number *C*_*l*_ + *C*_*r*_ = −, − − {25, 32, 39, 46}. The value of − was drawn randomly on each trial. We imposed this constraint to ensure that subjects had to attend to the click sounds in both ears. Stimulus presentation was also subject to the constraint that two consecutive clicks had to be separated by at least 5 ms. Furthermore, during pilot testing, subjects often claimed that they were biased toward the ear that presented either the first or the last click. To avoid such possible bias, the first and the last clicks in each stimulus occurred in both ears simultaneously, at time 0.0 s and 1.0 s, respectively. Thus, each ear received at least 2 clicks, and at most − − 2 clicks. We generated ten random versions of each of the 130 possible combinations of *C*_*l*_ and *C*_*r*_, and loaded the corresponding files into the memory of the custom program prior to the start of each session.

### 2.4. Online adaptive procedure

We set the difficulty of the perceptual task such that subjects were correct in approximately 60% of the trials. We achieved this using an adaptive staircase procedure (Kubanek et al. 2013). This procedure allowed subjects to perform close to the desired accuracy (first block: 61.9 ± 2.8% (mean±s.d., *n* = 54); second block: 60.3 ± 8.7%).

### 2.5. Handedness score

When recruiting subjects, we encouraged the participation of left handed subjects, to obtain as balanced a proportion of right- and left-handed subjects as possible. After each subject completed testing, they answered a set of question that probed the subject’s handedness. We used a set of questions based on the revised Edinburgh Handedness Inventory (Williams 1986). This test returns a number between −100 (strongly left-handed) and +100 (strongly right-handed). The mean±s.d. score over our subjects was +41.2 ± 57.0. In Fig. 3, we divided the subjects into three groups based on the handedness score. Subjects with a score higher than +33 were considered right-handed, subjects with a score less than −33 left-handed, and the subjects with a score between −33 and +33 were considered ambidextrous.

## 3. Results

### 3.1. Choice Behavior

Fig. 2A shows subjects’ choice behavior when they could choose either response freely (during the 20% of control trials in which no stimulus was presented). As seen in the figure, on those trials, subjects showed a bias to choose the rightward key (congruent key assignment block: mean 58.3%, difference from 50%, *t*_53_ = 3.85, *p* = 0.00032, two-sided t-test; reversed key assignment block: 57.4%, *t*_52_ = 3.16 (one subject did not perform the reversed block), *p* = 0.0026). Because there was no stimulus, there should be no difference between the choice proportions in the two blocks. Indeed, these proportions were statistically indistinguishable (*t*_52_ = 0.29, n.s., paired two-sided t-test).

**Fig. 2.**
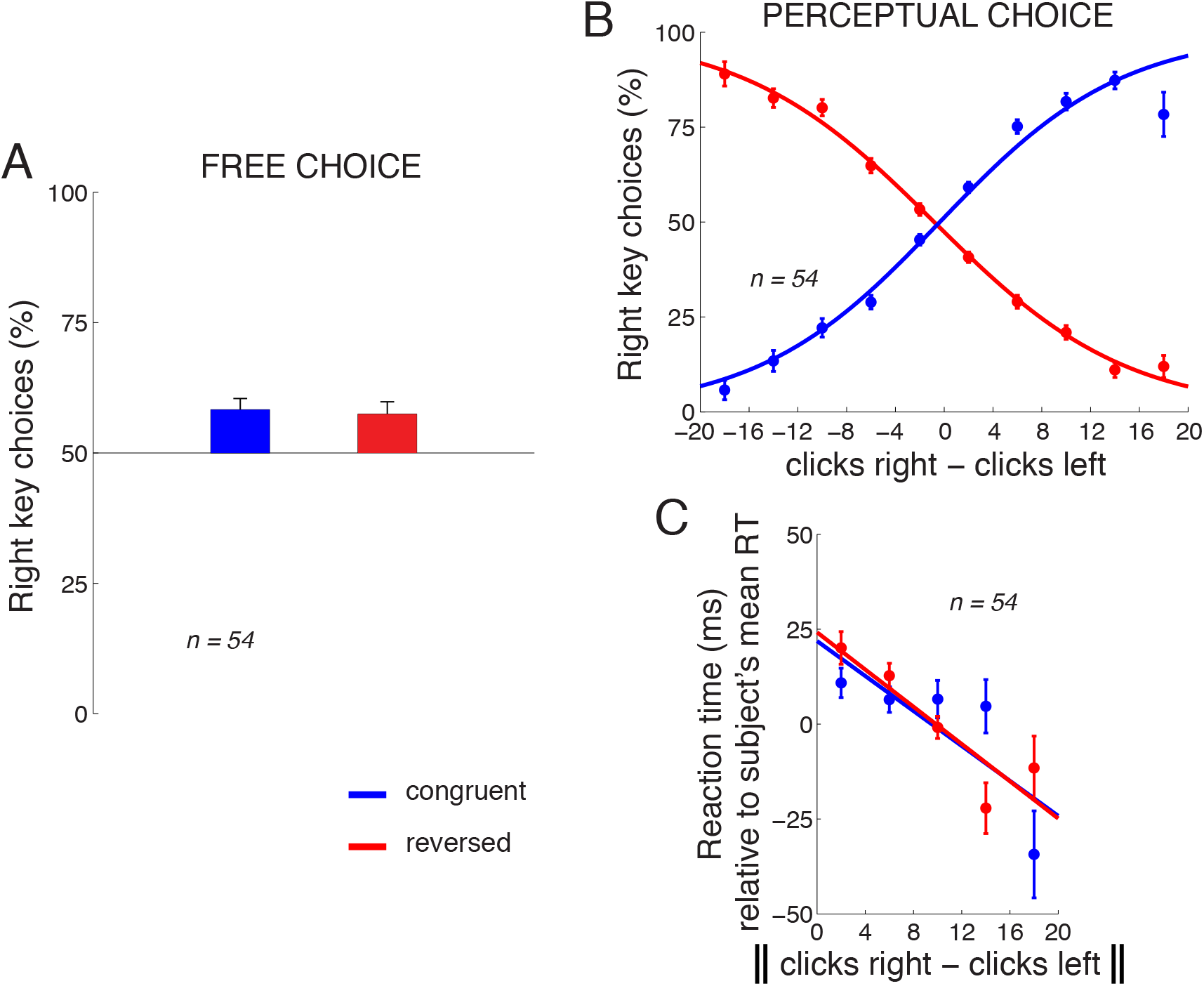
Choice behavior. A) Mean *±* s.e.m. proportion of rightward choices in the trials in which no stimulus was present and subjects chose freely. The data are shown separately for the congruent (blue) and reversed (red) response assignments. B) Mean *±* s.e.m. proportion of rightward choices as a function of the difference in the number of clicks in the right and the left ear, separately for the congruent and reversed response assignments. The curves represent logistic fits to the 10 data points in each block. C) Mean *±* s.e.m. RT as a function of the absolute difference in the number of clicks in the right and the left ear, separately for the congruent and the reversed block. To control for differences in mean RT over the subjects (445 *±* 123 ms, mean *±* s.d.), the mean RT was subtracted from each RT value in each subject. The line is a fit to the 5 data points in each block. Congruent block, *n* = 54, reversed block, *n* = 53 subjects.

When subjects made decisions based on the perceptual stimulus, their responses followed the given instruction (Fig. 2B). Specifically, in the congruent block (blue), when subjects heard substantially more (e.g., 10 more) clicks in the right ear than in the left ear, subjects predominantly pressed the right key, and vice versa. When the instructed key assignment reversed, the choice behavior accordingly reversed (red). We quantified the choice behavior using logistic regression, as shown in the logistic fits in Fig. 2B.

We applied the logistic regression to the choice data of each individual subject. To determine whether the stimulus was a significant factor in guiding the subjects’ responses, we measured the weight assigned to the click difference in this regression. This weight significantly differed from zero over the subjects (congruent block: mean weight 0.17 per click, *t*_53_ = 12.92, *p* < 0.0001, two-sided t-test; reversed block:

The amount of information in the stimulus may influence the time it takes subjects to produce a response, the reaction time (RT). We indeed found that the more information in the stimulus (greater difference in the number of the clicks between the two ears), the faster the subjects responded (Fig. 2C).

We quantified this relationship by fitting a line to this relationship in each subject, and measured the slope of the line. The mean slope over the subjects in the congruent block was −1.74 ms per click, and this slope significantly differed from zero (*t*_53_ = −2.61, *p* = 0.012, two-sided t-test). In the reversed block, the mean slope was −2.85 ms per click (*t*_52_ = −3.99, *p* = 0.00021). Note that these numbers are averages of the slopes computed separately in each subject, compared to the slopes shown in Fig. 2C, which represent, for visualization purposes, fits to data combined across all subjects.

### 3.2. Effect of Handedness

We quantified the effects of handedness and the instructed stimulus-key assignment (“key assignment”) on choice using an ANCOVA model. In this linear model, the output variable is the proportion of choices of the rightward key, computed separately for each subject. The input factors are handedness (a number between −100 and +100, see Methods), and the key assignment (binary variable, congruent or reversed). The ANCOVA model assesses whether these two factors explain a significant portion of the variance in the proportion of the rightward choices.

We first investigated the effects of handedness and the key assignment during free choice, i.e., in the 20% of trials in which there was no stimulus and subjects could freely choose to press either the left or the right key. In these trials, the key assignment should have no effect on subjects’ choices because in these trials there was no stimulus and thus no stimulus-response association. Indeed, the ANCOVA model revealed no significant effect of key assignment on the proportion of rightward choices (*F*_1,103_ = 0.07; n.s., see also Fig. 2A).

Interestingly, the ANCOVA revealed that subjects’ choices were significantly impacted by their handedness in this task (*F*_1,103_ = 7.50, *p* = 0.0073). As expected in this stimulus-free task, there was no interaction between handedness and key assignment (*F*_1,103_ = 0.28, n.s.). We therefore averaged the choice proportions for each subject across the two blocks. We then plotted the average proportion of rightward choices as a function of handedness, separately for each subject (Fig. 3A). The figure reveals that the proportion of rightward choices increases with the extent to which the subject is right-handed. When these data are fitted with a line, the slope of the line indicates a 15.0% change in the percentage of rightward choices over the range of handedness, and this slope significantly differs from zero (*t*_52_ = 2.35, *p* = 0.023, two-sided t-test). When the two most extreme individuals are excluded (the individual with the lowest and the individual with the highest proportion of rightward choices), the slope reveals a 20.0% change in the percentage of rightward choices over the range of handedness, and this slope significantly differs from zero (*t*_50_ = 3.72, *p* = 0.00051, two-sided t-test).

**Fig. 3.**
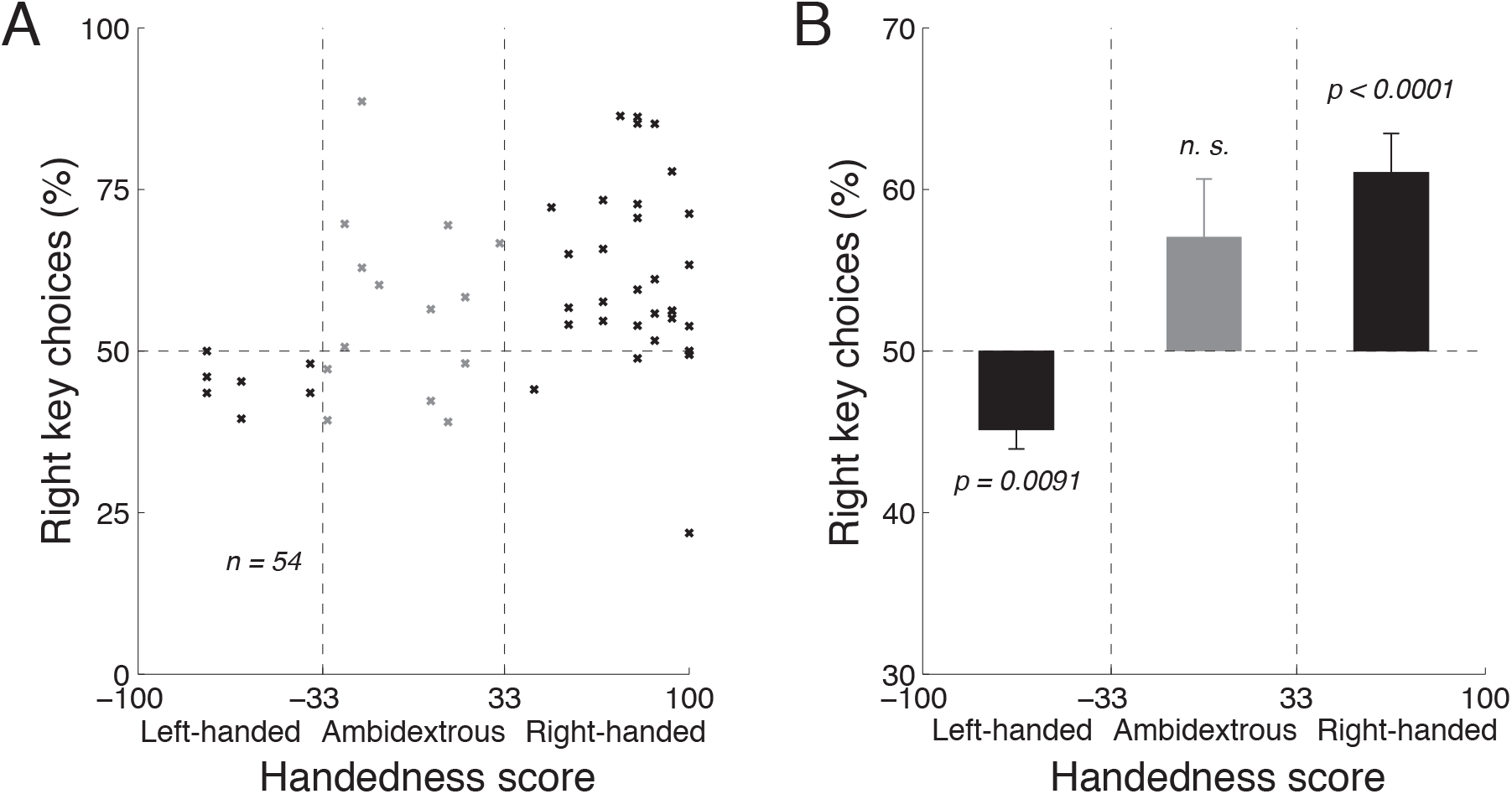
Handedness affects free choice. A) Mean proportion of rightward choices as a function of each subject’s handedness, in the trials in which there was no stimulus and so in which subjects chose freely. The vertical dashed lines segregate the effects in the left-handed, right-handed, and ambidextrous groups of subjects. B) The mean *±* s.e.m. effects computed using the data in A, separately for the left-handed, right-handed, and ambidextrous subjects. The *p* values give the significance of the test that a given mean proportion of choices significantly differs from the balanced 50% proportion (two-sided t-tests). In both A and B, data were pooled across the two blocks in each subject; *n* = 54 subjects.

We quantified the effects of handedness on choice separately for left-handed, ambidextrous, and right-handed subjects (Fig. 3B). Left-handed subjects (handedness scores lower than −33) preferred to make leftward choices (mean proportion of rightward choices, 45.2%), and this preference significantly differed from the balanced 50% (*t*_6_ = −3.82, *p* = 0.0088, two-sided t-test). In contrast, right-handed subjects (handedness scores higher than +33) strongly preferred to make rightward choices (mean proportion of rightward choices, 61.1%, *t*_32_ = 4.64, *p <* 0.0001). Ambidextrous subjects (handedness scores between −33 and +33) showed a tendency to choose the right key (mean proportion of rightward choices, 56.8%), but this tendency was not significantly different from the balanced 50% (*t*_13_ = 1.81, *p* = 0.093). Thus, when subjects make a choice based on their own deliberation, handedness is a significant factor in guiding the choice. It is surprising to observe an effect of handedness during choice that involves a movement as trivial as pressing a key with an index finger.

### 3.3. Effect of Auditory Processing Asymmetry

Finally, we tested whether and how handedness and the key assignment affect choice during perceptual decisions, i.e., in the trials in which subjects’ choices were guided by the auditory stimulus (Fig. 2B top). In contrast to the free choice, when subjects’ choices were based on the perceptual stimulus, the ANCOVA did not reveal an effect of handedness on choice (*F*_1,103_ = 0.04, n.s.). The potential effect of handedness was not masked by a difference across the key assignment blocks, because the ANCOVA also did not detect an interaction between handedness and key assignment (*F*_1,103_ = 0.3, n.s.). In contrast, in this task, the ANCOVA revealed a highly significant effect of the key assignment on choice (*F*_1,103_ = 20.76, *p <* 0.0001).

The effect of the key assignment on choice is shown in Fig. 4. The figure reveals that in the congruent block of trials, subjects preferred to make, on average, a rightward choice (blue). This effect (mean 53.2%) significantly differs from 50% (*t*_53_ = 2.94, *p* = 0.0049, two-sided t-test). This rightward choice preference could be either due to a movement-related effect or a sensory-related effect. In particular, in the congruent block, the effect may indicate an enhanced representation of the motor plan to press the right index finger, but it may also reflect an enhanced representation of the number of click sounds presented to the right ear. Reversing the key assignment provides a means to distinguish between these two possibilities. Intriguingly, when the key assignment reversed, the subjects’ choice preference also reversed (red). Subjects now preferred to choose the leftward option (mean rightward choices, 46.6%) and this leftward choice bias was significant (*t*_52_ = −3.64, *p* = 0.00063). The finding of a reversal of the rightward preference upon the reversal of the key assignment rules out a general rightward response bias. Instead, the effect indicates an enhanced representation of the click sounds presented to the right ear.

**Fig. 4.**
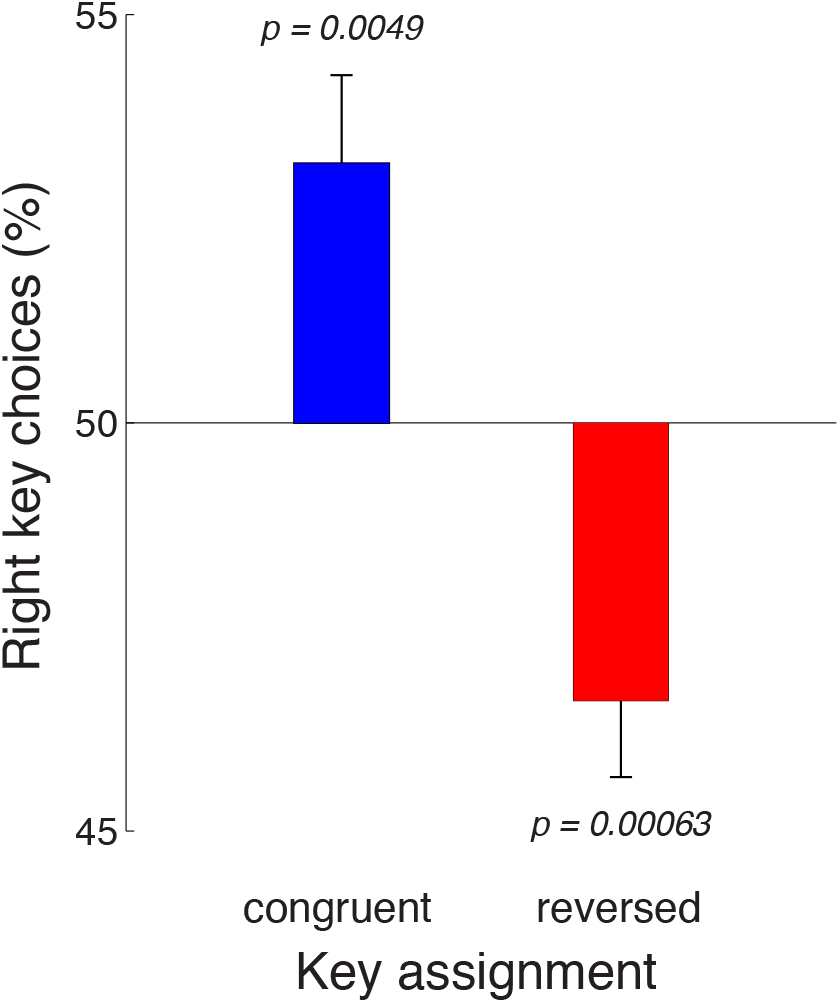
Auditory decision-making is affected by a right-ear enhancement. Mean *±* s.e.m., over the individual subjects, proportion of rightward choices in the task in which subjects’ choices were based on the auditory stimulus. The data are shown separately for the congruent (blue) and reversed (red) key assignments. The rightward choice preference (blue) reverses when the key assignment is reversed (red). This indicates an enhancement of the representation of sensory evidence presented to the the right ear. Congruent block, *n* = 54, reversed block, *n* = 53.

Handedness did not substantially impact this effect; as mentioned above, there was no interaction between the key assignment and handedness. Furthermore, the ANCOVA found no effect of or interaction with handedness (*p >* 0.55) when we only considered stimuli with balanced number of clicks in the right and the left ears (absolute difference in clicks less than two).

## 4. Discussion

Traditional models of information processing have held the view that higher-order cognitive processes, such as decision-making, function independently of bodily influences (Markman and Dietrich 2000; Wilson 2002; Clark 1999). These models have over the past several decades dominated cognitive science (Markman and Dietrich 2000; Wilson 2002; Clark 1999), artificial intelligence (Barsalou 2008; Ghazanfar and Turesson 2008), behavioral economics (Tversky and Kahneman 1981), and systems neuroscience (Schall 2002; Gottlieb 2007; Padoa-Schioppa and Assad 2008). According to these models, sensory information is passed to and processed by a centralized cognitive processing module. The outcome computed by this module, such as a decision, is subsequently passed on to the motor system to be transformed into a desired action.

This view has recently been challenged. Work on embodied cognition suggests that higher-order cognitive processes may be influenced by physical properties of the body and aspects of motor and sensory systems, as well as the computations in these systems (e.g., Abrams and Balota (1991); Clark (1998, 1999); Pfeifer and Bongard (2006); Barsalou (2008); Ghazanfar and Turesson (2008); Thelen et al. (2001)). In particular, experiments have demonstrated that many seemingly abstract cognitive processes—such as, interpreting emotion (Niedenthal 2007), processing language (Fischer and Zwaan 2008) and numbers (Domahs et al. 2010), memory retrieval (Dijkstra et al. 2007), or cognitive control (Brockmole et al. 2013; Weidler and Abrams 2014)—may be tightly associated with sensorimotor elements of the body.

We now show that the process of making a simple perceptual decision is also embodied. Specifically, we found that subjects’ choices depend on motor- and sensory-related attributes of the decision-maker—the subject’s handedness, and biases in the auditory perceptual system. Handedness substantially influenced subjects’ choices on trials in which subjects deliberately chose whether to press a left key with a left finger or a right key with a right finger. On trials in which the choices were based on a stereo auditory stimulus, subjects’ choices revealed a right ear-related enhancement and no handedness bias. We discuss the significance of these findings below.

It has been shown that the selection of which hand to use to reach for an object is influenced by subjects’ handedness (Bryden et al. 1994; Bishop et al. 1996; Gabbard et al. 1998; Calvert 1998; Stins et al. 2001). Specifically, in these tasks and their variants, an experimenter systematically varies a particular property or location of an object within a subject’s workspace. Subjects are then asked to reach for or manipulate the object using the hand of choice. It is commonly found that the relative frequency with which a hand is selected is a function of handedness: right-handed subjects on average prefer to reach for an object using the right hand, whereas left-handed subjects prefer to reach for an object with the left hand. In these studies, a reach for, a grasp of, or a manipulation of an object involves a relatively complex movement that requires dexterity and often further engages a higher-order computation that must weigh which hand is more suitable for or efficient in successfully completing the movement (Bryden et al. 1994; Bishop et al. 1996; Gabbard et al. 1998; Calvert 1998; Stins et al. 2001). It is then perhaps unsurprising that to successfully perform such movements, subjects prefer to use the hand that they have been using predominantly for such purpose throughout their life (Serrien et al. 2006).

In contrast to such complex movements, in our study subjects performed a simple key press using a finger. The left index finger was positioned over one key and the right index finger over another. In this state, subjects decided which of the keys to press. Left-handed subjects showed a bias in pressing the left key, whereas right-handed subjects showed the opposite bias (Fig. 3). Given the simplicity of the movement, it is surprising to find that subjects’ choices were affected by handedness. This suggests that the effect of handedness may extend beyond a choice of a dexterous movement; it may impact choices that involve a movement of the hand in general, even if such movement is trivial. This is supported by electrophysiological and interventional studies (Brasil-Neto et al. 1992; Kim et al. 1993; Stancák Jr and Pfurtscheller 1996; Schluter et al. 1998; Solodkin et al. 2001; Serrien et al. 2006) that report neural signatures of hand dominance in premotor and motor regions, regions that plan and execute both complex and simple movements.

Notably, the effect of handedness was observed in a task in which no stimulus was present. When the subjects’ choices were guided by the perceptual stimulus, there was no significant effect of handedness on choice (and no handedness-based interaction). This is in line with findings made in dichotic word discrimination tasks in which handedness had no or minimal effect on subjects’ judgements (Curry 1974; Kimura et al. 1983). Thus, the influence of handedness on choice may be suppressed when a decision is guided by sensory information. Our data reveal that the prominent effect of handedness on choice is observed specifically during *self-initiated* choices.

When the subjects’ choices were based on the stereo auditory stimulus, subjects showed a significant rightward choice bias with the congruent key assignment (Fig. 4, blue). When the key assignment reversed, the effect reversed to a significant leftward bias (Fig. 4, red). These two effects together indicate that there is an enhancement in the perception of the sensory information presented to the right ear. In this regard, there are findings of asymmetric representations of auditory information in the literature. In particular, in dichotic listening tasks, each ear is simultaneously presented with spoken words. In these tasks, subjects correctly identify more words presented to the right ear compared to words presented to the left ear (Kimura 1961a,b; Broadbent and Gregory 1964). However, interestingly, this effect is specific to verbal material; sounds that are not words, such as vocal and non-vocal environmental sounds, appear to show a weak reverse effect—a left ear superiority (Knox and Kimura 1970; King and Kimura 1972). The presence of a right ear advantage specifically in verbal tasks has led to the suggestion that the effect may reflect the lateralization of language processing to the left hemisphere, given the dominance of the crossed auditory pathways over the uncrossed pathways (Geschwind and Galaburda 1984; Kimura 2011). We found a right-ear advantage in a non-verbal task, a task that requires an accumulation of discrete quanta of auditory evidence over time (Brunton et al. 2013). Because the stimuli were non-verbal, the effect we report may have a different origin than the word-specific right ear advantage reported previously (Kimura 2011). It will be important to determine at which stage of the sensorimotor transformation the effect occurs.

The effect could act on early sensory representations (biasing the representation of the auditory evidence), but could also act further downstream (enhancing a “rightward” decision). One way to disambiguate between these possibilities is to obtain evidence from other sensory modalities, e.g., visual. If the same effects of presentation side were observed in the visual task, that would suggest that there is a bias that generally favors “rightward” decisions. However, to our knowledge, no such general “rightward” bias has been suggested in the decision literatures.

In summary, we found that in a simple perceptual decision task, subjects’ choices were influenced by their handedness and an enhancement of perceptual evidence presented to the right ear. When subjects were free to choose to make a simple finger movement with either hand, their choices were biased by handedness. When their decisions were based on a stereo auditory stimulus, the choices indicated a bias towards the right ear, and no effect of handedness. Thus, the seemingly deliberate process of making a simple choice can be partially embodied, skewed by asymmetries of the human motor and sensory systems.

## Acknowledgments

This study was supported by the NIH grants EY012135 and EY002687. The authors declare no competing conflict of interest.

## Notes

### Competing Interest Statement

The authors have declared no competing interest.

### Summary of Updates

Fixed several typoes in the Abstract. Did not edit anythign else.

